# Lack of functional polyester-biodegrading potential in marine *versus* terrestrial environments evidenced by an innovative airbrushing technique

**DOI:** 10.1101/2024.11.06.622231

**Authors:** Alberto Contreras-Moll, Theo Obrador-Viel, Rocío Daniela Inés Molina, Maria del Mar Aguiló-Ferretjans, Balbina Nogales, Rafael Bosch, Joseph A. Christie-Oleza

## Abstract

Biodegradable plastics, primarily aliphatic polyesters, degrade to varying extents in different environments. However, the absence of easily implementable techniques for screening microbial biodegradation potential —coupled with the limitations of non-functional omics analyses— has restricted comparative studies across diverse polymer types and ecosystems. In this study, we optimized a novel airbrushing method that facilitates functional analyses by simplifying the preparation of polyester-coated plates for biodegradation screening. By repurposing an airbrush kit, polyester microparticles (MPs) could be evenly sprayed onto solid media, enabling rapid detection of extracellular depolymerizing activity via clearing zone halos. This technique was effective in screening both isolated microbial cultures and natural environmental samples, demonstrating its versatility. The method was successfully applied across multiple environments, ranking the biodegradability of six polyesters, from most to least biodegradable: polycaprolactone (PCL), poly[(R)-3-hydroxybutyrate] (PHB), poly(butylene succinate) (PBS), poly(ethylene succinate) (PES), poly(lactic acid) (PLA), and poly(butylene adipate-co-terephthalate) (PBAT). Most notably, it revealed a consistent 1,000-fold higher biodegradation potential in terrestrial compared to marine environments. This approach offers a valuable tool for isolating novel polyester-degrading microbes with significant biotechnological potential, paving the way for improved plastic waste management solutions.

**SYNOPSIS:** Screening polyester-degrading microbes has traditionally been challenging due to the difficulty of incorporating polyesters into solid media. This study introduces an innovative airbrush spraying method that simplifies the preparation of polyester MPs on solid media plates, making the screening process significantly more efficient. Using this method, we demonstrated the differential biodegradability of aliphatic polyesters across environments. The results revealed a stark contrast in biodegrading potential, with terrestrial ecosystems exhibiting substantially higher activity compared to marine environments.

**Abstract graphic:** 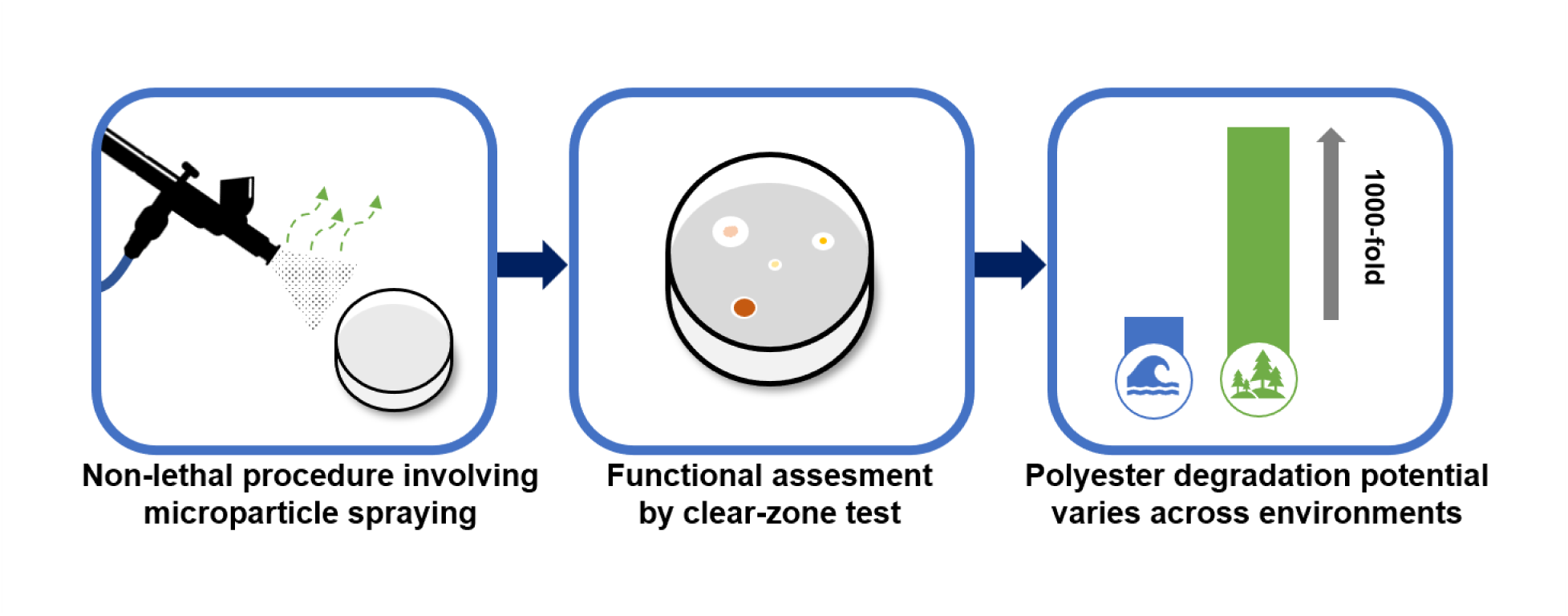

## INTRODUCTION

The intrinsic biodegradability of polyesters stems from the presence of ester bonds (-C(=O)-O-C-) in their structural backbone, in contrast to traditional plastics, which possess highly inert chemical structures based on -C-C-bonds.^1^ Hence, biodegradable plastics -mainly polyesters-arose as an eco-friendly promising alternative to traditional plastics, aiming to alleviate plastic accumulation in the environment thanks to their biodegradability.^2–3^ Polyester biodegradation is mainly determined by the interplay between: *i*) environmental conditions (temperature, pH, moisture and C/N ratio), which directly control hydrolysis reaction rates and vary greatly between environments, and *ii*) the material’s physicochemical properties, (crystallinity, molecular weight and chain flexibility), which differ between polyester types and may hinder biodegradation.^4–7^ On top of the aforementioned factors, efficient polyester biodegradation in the environment also relies on the presence and recruitment of microorganisms able to hydrolyze and assimilate such materials.^8,9^ The widespread taxonomic diversity of microbes capable of degrading polyester, along with the vast array of associated hydrolases, complicates the assessment of true biodegradation potential within microbiomes using non-functional omics techniques.^10–12^

Functional evaluation using traditional culture-dependent techniques have been used to obtain biotechnologically-relevant microbial isolates -and their associated enzymes-capable of degrading different polyesters.^11,13–16^ To perform the screening of promising degrading strains from environmental samples, the inoculum is normally grown on a solid mineral medium containing the polyester as the only carbon source, which is homogenized as effectively as possible in order to generate an opaque plate. Then, the selection of biodegrading microbes is based on the appearance of a clearing halo around colonies. This approach is known as the “clear zone test”, where hydrolysis halos are formed as a result of the polyester depolymerization by extracellular hydrolases secreted by the cells.^17^ To date, there are a number of studies that have used this method to isolate microbes that can hydrolyze polyesters like PBAT, PBS, PCL, PES, PHB and PLA.^18–25^ However, since polyester-based plastics are practically insoluble in aqueous solutions, the preparation of plates is usually preceded by a laborious emulsification protocol to successfully homogenize the plastic in the medium before solidifying, altogether not guaranteeing success.^19,24^ The complexity and considerable costs associated with the emulsification procedure hinder the efficient screening of polyester degraders in the environment -particularly for plastics like PBAT or PLA- and limit our understanding of plastic biodegradation dynamics across natural habitats. This situation recently led to the design by Shin et al. (2021) of an ingenious alternative to incorporate the plastic in the medium in the form of air-blown MPs.^26^ Their work has resurrected the concept of “air-spraying” -originally presented by Kiyohara et al. (1982) for water-insoluble solid hydrocarbons-as a means to distribute hydrophobic substrates on solid agar-containing medium.^27^ Briefly, the polyester is dissolved in an organic solvent -preferably one with high volatility- and directly sprayed onto an inoculated agar plate with the use of a customized spraying system. This approach allows a rapid identification of polyester-degrading microorganisms, effectively reduces the operational time, avoids the addition of carbon-based emulsifying agents and guarantees a successful preparation of plates.^26^ However, user accessibility was neglected and scarce details about the intricate spraying system were provided, altogether preventing an easy implementation into routine laboratory workflows.

In view of the promise of the air-spaying technique, we have refined and optimized this innovative method by repurposing an airbrush kit as an affordable and accessible spraying system that can be easily implemented in any laboratory. A comparison with the emulsification method and other technical aspects such as procedural toxicity and the particle size were evaluated in detail. After optimization on both microbial isolates and environmental samples, the method was used to determine the plastic-biodegrading potential and frequencies of polyester-hydrolyzing microorganisms across multiple environments. To facilitate widespread adoption, we conclude by presenting a detailed user-friendly protocol, supporting the establishment of this method as a robust approach for isolating potential plastic-biodegrading microorganisms.

## MATERIALS AND METHODS

### Bacterial strains and culture conditions

All microbial strains used in this work were routinely grown on Luria-Bertani (LB) agar plates before the commencement of the experiments. These strains include *Amycolatopsis orientalis* DSM 40040^T^ and *Thermobifida fusca* DSM 43793, obtained from the German Collection of Microorganisms and Cell Cultures (DSMZ) and used as positive controls to test the formation of hydrolysis halos in PLA- and PBAT-sprayed plates, respectively^17,28^; and soil-dwelling isolates *Pseudomonas* sp. J10SM and *Rhodococcus* sp. J04LL together with *Kodamaea* sp. A6, previously obtained from insect guts. Bushnell-Haas (BH) basal mineral medium was used to conduct all experiments involving polyester hydrolysis.^29^ pH was adjusted to 7.0 and yeast extract (0.001 % w/v) was added as a source of vitamins and growth factors upon autoclaving. Agarose 0.5 % w/v (Condalab, Spain) was designated as the solidifying agent to facilitate microbial growth and the discrimination of hydrolysis halos. Agar (Condalab, Spain) 1.5 % w/v was used for the screening of polyester degraders in environmental samples as a more affordable alternative when preparing large volumes of plates. BH medium was supplemented with 3 % w/v NaCl when working with samples of marine origin. A mix of 0.1 % w/v labile carbon sources including pyruvate, succinate, glucose and glycerol was used to determine the total colony forming units (CFU) in environmental samples. Extracellular proteolysis potential was investigated by preparing BH agar plates with 1.0 % w/v skim milk powder (Nestlé, Spain). Emulsified PCL plates were prepared by dissolving 0.1 g of PCL in 8 mL of acetone, after which it was added to 100 mL of temperate BH agarose medium, shaken and poured in Petri dishes.^24^ Remaining acetone in the plates was left to evaporate at 30 °C for 24 hours before use. It is important to note that only PCL plates were prepared in this way, and no success was achieved with other polyesters. Polyester spray-coated plates are detailed below.

### Polymer dissolution

Five aliphatic polyesters (PBS, PCL, PES, PHB and PLA) and an aromatic-aliphatic polyester (PBAT) (Table 1) were dissolved separately in the organic solvent dichloromethane (DCM; Labkem). Dissolutions were prepared in glass bottles containing 20 mL of DCM with a polyester concentration of either 1.5 % or 3.0 % w/v, depending on the plastic (Table 1). To accelerate polymer dissolution, glass bottles were placed in an ultrasound bath (J.P. Selecta S.A., Spain) and subjected to external sonication at 50-60 Hz for varying amounts of time (Table 1). PHB required additional mechanical stirring for 30 minutes to efficiently disaggregate and dissolve the powder in DCM.

**Table 1.**
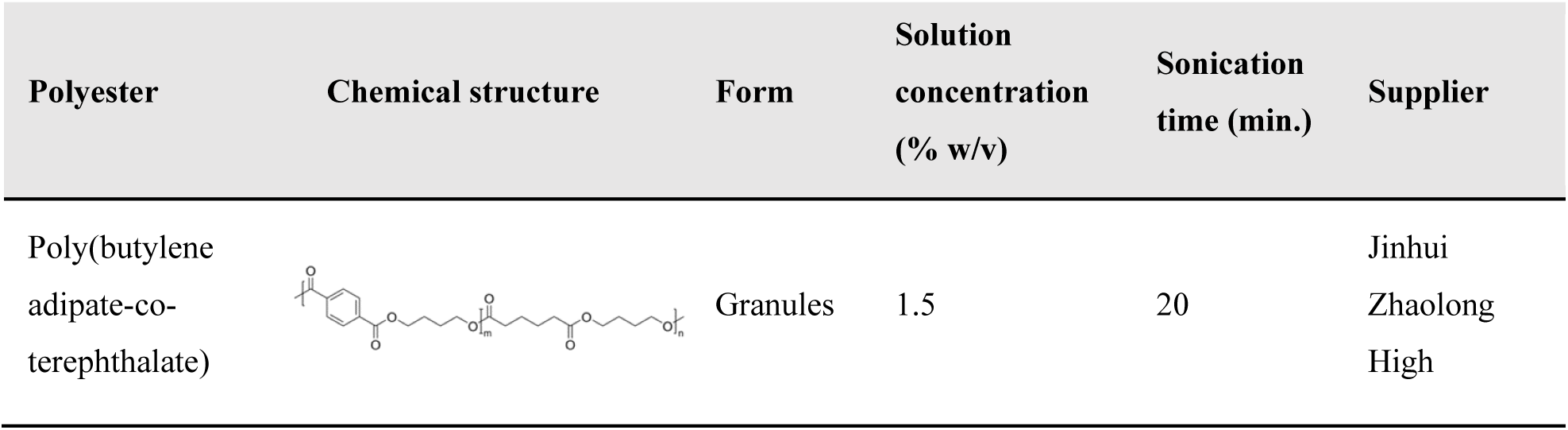
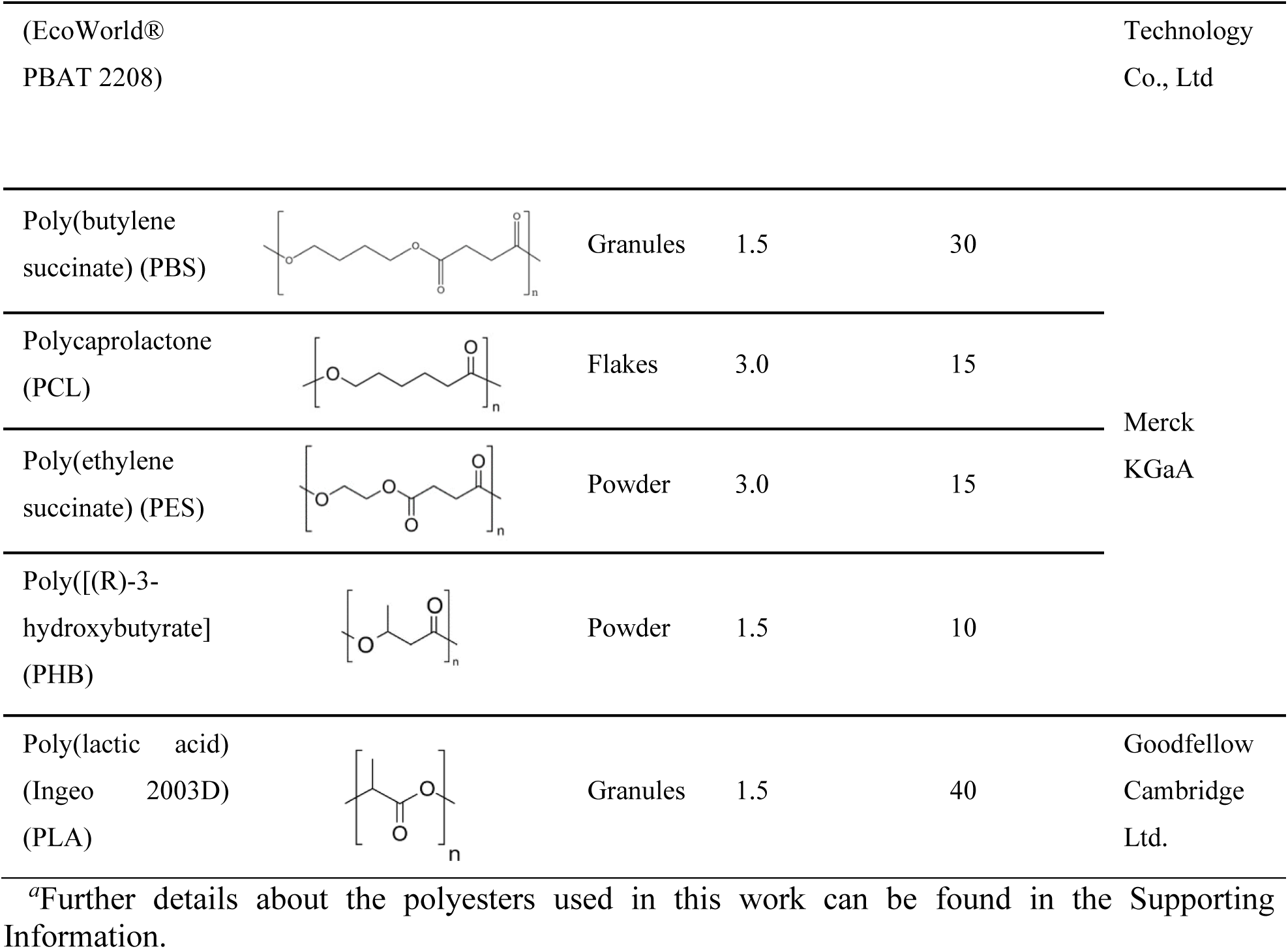
Polyester details and DCM dissolution preparation*^ɑ^*.

| Polyester | Chemical structure | Form | Solution concentration (% w/v) | Sonication time (min.) | Supplier |
| --- | --- | --- | --- | --- | --- |
| Poly(butylene adipate-co-terephthalate) |  | Granules | 1.5 | 20 | Jinhui<br>Zhaolong<br>High |
| (EcoWorld® PBAT 2208) |  |  |  |  | Technology Co., Ltd |
| Poly(butylene succinate) (PBS) |  | Granules | 1.5 | 30 | Merck KGaA |
| Polycaprolactone (PCL) |  | Flakes | 3.0 | 15 |  |
| Poly(ethylene succinate) (PES) |  | Powder | 3.0 | 15 |  |
| Poly([(R)-3-hydroxybutyrate] (PHB) |  | Powder | 1.5 | 10 |  |
| Poly(lactic acid) (Ingeo 2003D) (PLA) |  | Granules | 1.5 | 40 | Goodfellow Cambridge Ltd. |

<sup>a</sup>Further details about the polyesters used in this work can be found in the Supporting Information.

### Polyesters spray coating

A commercially available airbrush starter kit consisting of a FE-130 gravity-fed airbrush coupled to an FD18-2K air compressor (Fengda®) was used to spray polyester dissolutions directly onto readily inoculated translucent solid media plates in a fume hood (Figure S1). Briefly, 1 mL of each plastic dissolution was sprayed through a 0.5 mm nozzle at a constant pressure of 3 bars (measured at the air compressor pressure gauge) onto the agarose plates. Plastic was sprayed in a circular motion whilst keeping a working distance of 10-15 cm between the airbrush nozzle and the solid plate surface (Figure S1). After spraying, Petri plates were covered and left untouched in the fume hood for 30 minutes at room temperature to let any remaining solvent traces evaporate, after which they were incubated for optimal microbial growth. Hydrolysis halos were examined by plain sight as well as with the aid of a stereoscopic microscope (Motic® SMZ-168 series) to further verify particle hydrolysis. A more detailed protocol is included as Supporting Information.

### Polyester particle size calculation and morphology assessment

Freshly sprayed plates were inspected to assess particle morphology and size distribution in terms of their diameter upon solvent evaporation. Optical microscope images were taken for each polyester type using a Moticam 5.0 MP digital camera coupled to a Motic® BA310LED optical microscope (150x magnification). A total amount of 440 particles were randomly selected per polyester and their surface area calculated using ImageJ.^30^ Particle diameter was derived from the raw data to construct the final particle size distribution (median, minimum and maximum values) using Prism (v8.0.2, GraphPad). For comparison, PCL emulsions were prepared by dissolving 0.1 g of PCL in 8 mL of acetone, which was then poured into 100 mL of distilled water and stirred at 80 °C for several hours to evaporate the organic solvent. Particles formed in PCL emulsions were then analyzed by static laser light scattering (SLS) using a Microtrac SYNC particle size and shape analyzer and the size distribution was obtained with 5 mL of the original emulsion. Complementarily, scanning electron microscopy (SEM, HITACHI S3400N) and high-resolution transmission electron microscopy (HRSTEM, Talos F200i) images were obtained to provide a closer look at the morphology of the particles.

### Toxicity assays

Freshly grown colonies on LB were resuspended in Ringer solution by vortexing (OD_600_ 0.1) and serially diluted by 10-fold. Then, 10 µL from each dilution was inoculated by triplicate on LB agar plates. Two plates were inoculated for each strain: one of them was air-sprayed with a PCL solution in DCM, whereas the non-treated plate was used as control. Plates were placed in an incubator at 30 °C until colonies were visible. Colony counting was performed using the dilutions that yielded between 30 and 300 CFU.

### Screening of polyester degraders in a culture collection

Single colonies from our laboratory culture collection were picked and transferred to microcentrifuge tubes containing 0.5 mL of BH medium to prepare liquid inocula (OD_600_ 0.1). Then, 10 µL of each inoculum was spotted in triplicate on BH agarose plates and left to dry in a laminar flow cabinet before plate-spraying with each polyester. Coated plates were then incubated at either 20-30 °C (mesophiles) or 50 °C (thermophiles) and inspected for the appearance of clearance halos around the inoculation spots during the course of three weeks.

### Environmental sample collection

The screening of polyester biodegrading potential was carried out on samples collected from different locations and processed on the same day (Table 2). All soils and sediments were retrieved directly from the top 10 cm layer and kept in 50-mL Falcon tubes for transport to the laboratory. Sediments from the ports were collected using a sediment grab. Dry soils and composts were further sieved to remove large particles (> 1.5 mm) after which 1 g was added to 9 mL of BH medium, thoroughly vortexed at maximum intensity for 1 minute to effectively disaggregate soil particles and obtain a microbial suspension. Freshwater and marine sediment samples were processed in a similar way using the freshwater or seawater from the same location for microbial resuspension instead of BH medium. Prior to use, dense and buoyant debris was left to separate for 10 minutes. The microbial suspension was further used for plating as detailed below. Dry weight was calculated by subjecting the samples to air-drying in an oven at 60 °C before gravimetric analysis.

**Table 2.** Environmental samples origin.

| Environment | Sample | Content | Origin | Geographic location |
| --- | --- | --- | --- | --- |
| Compost | CoT | Commercial compost soil (Semillas Batlle, Spain) | - | - |
|  | CoTir | Industrial composting facility compost soil | Palma, Spain | - |
|  | CoA | Homemade compost soil | Palma, Spain | - |
| Soil | TV | Vineyard soil | Palma, Spain | 39°64'05.2"N<br>2°64'36.6"E |
|  | TA | Almond tree soil | Palma, Spain | 39°64'10.5"N<br>2°64'35.0"E |
|  | TO | Olive tree soil | Palma, Spain | 39°64'11.8"N<br>2°64'34.7"E |
| Freshwater sediment | ToE | Freshwater sediment (seasonal stream) | Esporles, Spain | 39°67'04.3"N<br>2°57'72.5"E |
|  | ToT | Freshwater sediment (spring) | Esporles, Spain | 39°66'46.7"N<br>2°57'15.3"E |
|  | ToSJ | Freshwater sediment (irrigation channel) | Palma, Spain | 39°32'44.5"N<br>2°45'03.1"E |
| Marine port sediment | PP | Marine sediment (port) | Palma, Spain | 39°56'81.2"N<br>2°63'88.7"E |
|  | PC | Marine sediment (port) | Palma, Spain | 39°56'13.2"N<br>2°66'76.3"E |
|  | PS | Marine sediment (port) | Llucmajor, Spain | 39°50'06.0"N<br>2°74'56.9"E |
| Marine coastal sediment | CS | Marine sediment (coast) | Llucmajor, Spain | 39°49'86.7"N<br>2°74'76.5"E |
|  | CA | Marine sediment (coast) | Palma, Spain | 39°52'93.2"N<br>2°72'88.6"E |
|  | CB | Marine sediment (coast) | Llucmajor, Spain | 39°48'70.3"N<br>2°73'66.0"E |

### Screening of polyester degraders in environmental samples

Aliquots of 2 mL were taken from each suspension and subjected to serial decimal dilution before plating 100 µL on BH agar plates (plates for marine samples were supplemented with 3 % w/v NaCl), and further spray-coated with the different polyesters. Duplicate plates were performed per dilution, polyester and sample. For proteolytic determinations, BH agar plates were supplemented with 1.0 % w/v skimmed milk and dilutions plated as indicated above. To perform the total CFU counting from each microbial suspension, 10 µL of each dilution was inoculated by triplicate droplets on BH agar plates containing the mix of labile carbon sources. Plates were incubated at 28 °C, and both colony and degradation halos were routinely counted over three weeks. The frequencies of polymer biodegraders were calculated by dividing the total number of halo-forming colonies by the total CFU grown on labile substrates.

### Spray *versus* emulsion method comparison

Aliquots were taken from the homemade compost soil suspension and subjected to serial decimal dilution after which 100 µL of each dilution was plated in triplicate on *i*) BH agarose plates that were later spray-coated with PCL, and *ii*) BH agarose plates containing emulsified PCL. Both total CFU and halo-forming colonies were counted visually and with the aid of a stereomicroscope after incubation at 28 °C for three weeks.

### Ribosomal DNA sequencing for microbial identification

Isolates showing the ability to hydrolyze at least one polyester were selected for 16S/18S ribosomal deoxyribonucleic acid (rDNA) sequencing and identification. Product amplification was conducted according to the colony polymerase chain reaction (PCR) method in a T100^TM^ Thermal Cycler (Bio-Rad, Germany).^31^ Samples were previously heated to 90 °C for 5 minutes before the addition of the PCR Master Mix and MyTaq polymerase (MyTaq^TM^ Mix, Bioline). For 16S rDNA amplification, primers 27F (5’ -AGAGTTTGATCMTGGCTCAG- 3’) and 1492R (5’ - TACGGYTACCTTGTTACGACTT- 3’) were used.^32^ 18S rDNA amplification was performed with primers 63F (5’-ACGCTTGTCTCAAAGATTA-3’) and 1818R (5’- ACGGAAACCTTGTTACGA- 3’).^33^ PCR amplification products were purified and sent to Macrogen, Inc. (Spain) for Sanger sequencing. Sequences were analyzed and trimmed using BioEdit v7.2.5 and compared against NCBI rRNA/ITS databases using BLASTn to perform the identification^34,35^

### Statistical analysis

Descriptive and statistical analyses were performed using GraphPad Prism v. 8.0.2 software. Normality of the data and homoscedasticity were analyzed by means of the Shapiro-Wilk test and F-test, respectively. Differences between groups were evaluated using the Student’s unpaired t test. A level of statistical significance of P < 0.05 was assumed for all tests.

## RESULTS AND DISCUSSION

### Sprayed plates: macro- and microscopic appearance

Solid media plates sprayed with polyester dissolutions showed an opaque appearance (Figure 1), a requirement for the phenotypic identification of polymer hydrolysis during the clear zone test. Opacity increased with the amount of plastic sprayed on the plate, which ranged from 15 mg to 45 mg (Figure S2). The quantities were optimized to 15 mg for PBAT, PBS, PHB and PLA, whereas it was increased to 30 mg for PCL and PES. This decision was based on the cost-effectiveness of the method: the amount of sprayed plastic was the minimum required to easily visualize halo-forming colonies. This is evident when comparing PHB-sprayed plates, which showed the opaquest and roughest surface with only 15 mg, as opposed to PCL- and PES-sprayed plates, which displayed a smoother and lighter appearance even with higher amounts of plastic (30 mg; Figure 1 and S2). These differences in appearance might stem from the interplay between the intermolecular forces established 1) amongst polymer chains and 2) between these chains and the solvent molecules during plate spraying. The outcome of these interactions is what dictates their solubilization in the solvent and, later, the particle formation during spraying.^36^ Considering the stock concentrations for each polyester dissolution (Table 1), the volume of dissolved polyester sprayed per plate was set to 1 mL for standardization between all materials. This volume was sprayed during a 4-second frame window under the conditions described in the methods section, providing enough time to easily distribute the polyester across the entire agarose plate. Similarly, the working distance between the plate and spraying nozzle was set at 10-15 cm so the solvent from the droplets would evaporate and solid polyester particles could form before reaching the agarose surface. Otherwise, droplets with remains of solvent would aggregate and merge into an undesirable plastic film when impacting on the plate.^37^

**Figure 1.**
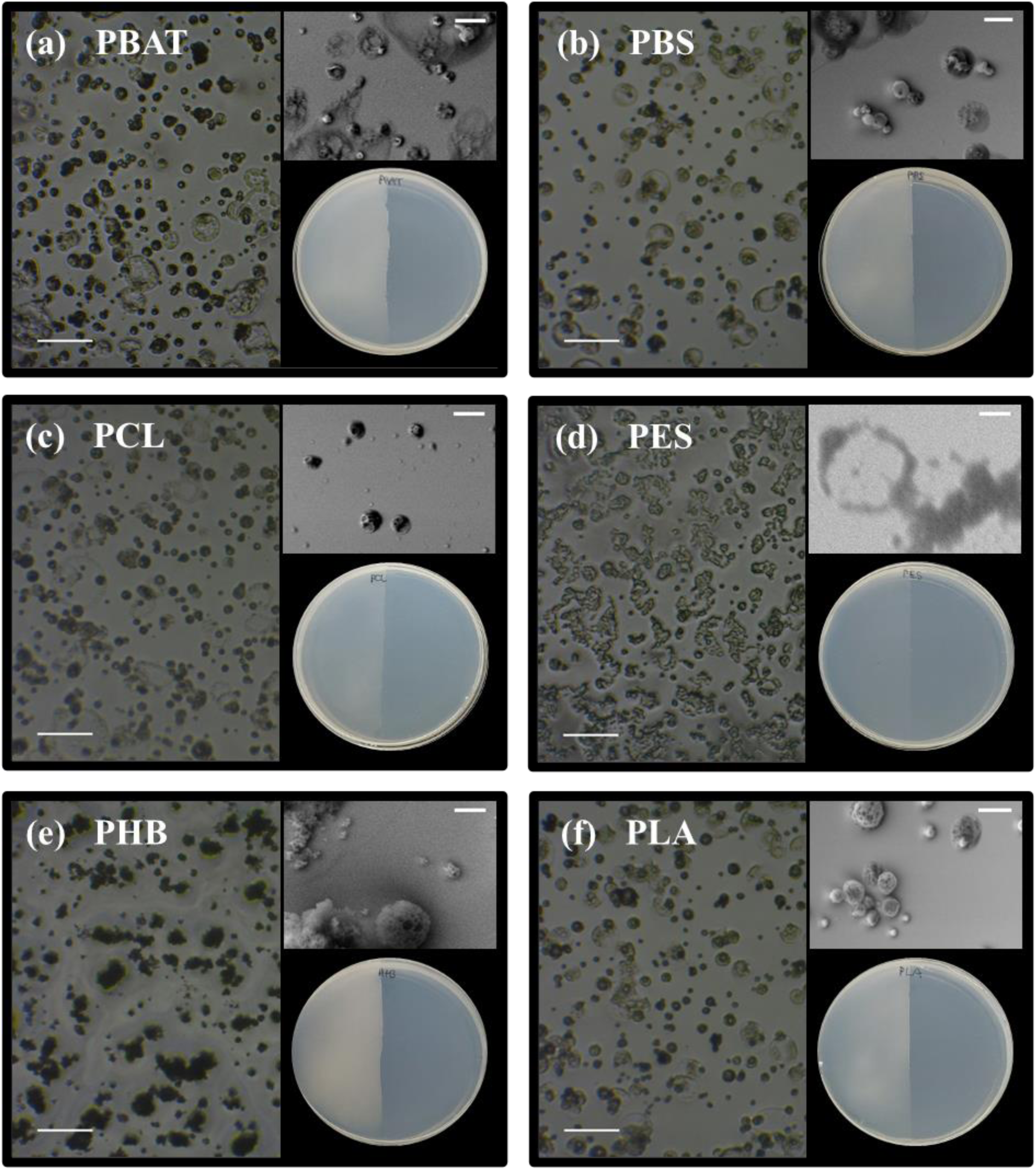
Polyester microparticles formed during the spraying method. Images were taken by light microscopy (150x magnification; scale bar: 100 µm) and SEM (top right corner of each image; scale bar: 20 µm). Left half spray-coated plates are included for visual comparison with non-coated right half (bottom right corner of each image). PBAT (a), PBS (b), PCL (c), PES (d), PHB (e), PLA (f).

Plates sprayed with the different polyesters were observed under a phase-contrast microscope to assess the size and morphology particles adopted during spraying (Figure 1). Polyesters scattered over the surface in the form of amorphous MPs with an overall median diameter of 14 µm (Figure S3). While 50% of the MPs possessed a diameter between 10 µm and 20 µm, MPs of 2.6 µm and up to 78.2 µm were observed (Table S1). Regarding their morphology, MPs were present in different shapes that varied depending on the polyester: 1) MPs with a spherical appearance, which were the most prominent amongst PBAT, PBS, PCL and PLA samples; 2) clearer flat plastic deposits, which dominated the PES particle population and accounted for the larger particulates in the rest of groups; and 3) granular MPs found in PHB sprayed plates (Figure 1). Hence, in most cases (PBAT, PBS, PCL, PHB and PLA), solid particles fully formed before reaching the agarose surface, giving rise to spherical MPs. It is important to note that MPs had pores in their structure, which has been reported as a secondary effect of solvent evaporation during spraying for highly volatile solvents, such as DCM.^38,39^ On the other hand, some droplets reached the surface before the solvent evaporated, forming larger clear flat deposits. This was especially obvious for PES sprayed plates (Figure 1d), for which the MP diameter was not measured due to the merge of deposits. PHB dissolutions in DCM showed a white appearance probably due to an uncomplete solubilization in the solvent. As a result, an unstable colloid of PHB granules, rather than a true solution might have been obtained during plastic solution preparation. This would explain why granular MPs -formed by the reaggregation of non-solubilized PHB granules during spraying- rather than spherical MPs were observed (Figure 1e).

### Assessment of solvent toxicity on inoculum viability during plate spray-coating

While microbial inocula can be added to pre-sprayed polyester plates, i.e. by depositing the inoculum as drops on the polyester-coated plate surface, most screening applications for plastic-degrading microbes from environmental samples will require spreading the cell suspension on the plate before applying the polyester. DCM is recognized as a toxic and irritating agent and potentially carcinogenic substance.^40^ When it comes to bacterial and fungal models, its toxicity resides mainly in its capability of modifying the membrane’s fluidity, increasing membrane permeability leading to cell death.^41,42^ Therefore, we examined whether the DCM dissolved polyester could affect cell viability on pre-inoculated plates. For this, three representative strains (i.e. gram-negative *Pseudomonas* sp. J10SM, gram-positive *Rhodococcus* sp. J04LL, and yeast *Kodamaea* sp. A6) were cultured on solid medium prior to spraying with DCM-dissolved PCL. No significant differences in cell viability or colony morphology were detected between the PCL-sprayed plates and non-sprayed controls (Figure 2): *Pseudomonas* sp. J10SM (control: 76.8 ± 9.9 CFU; PCL-sprayed: 72.8 ± 10.5 CFU; Student’s t-test; p = 0.5996), *Rhodococcus* sp. J04LL (control: 56.3 ± 7.4 CFU; PCL-sprayed: 54.5 ± 9.3 CFU; Student’s t-test; = 0.7135) and *Kodamaea* sp. A6 (control: 63.5 ± 4.7 CFU; PCL-sprayed: 64.0 ± 6.2 CFU; Student’s t-test; p = 0.9018).

**Figure 2.**
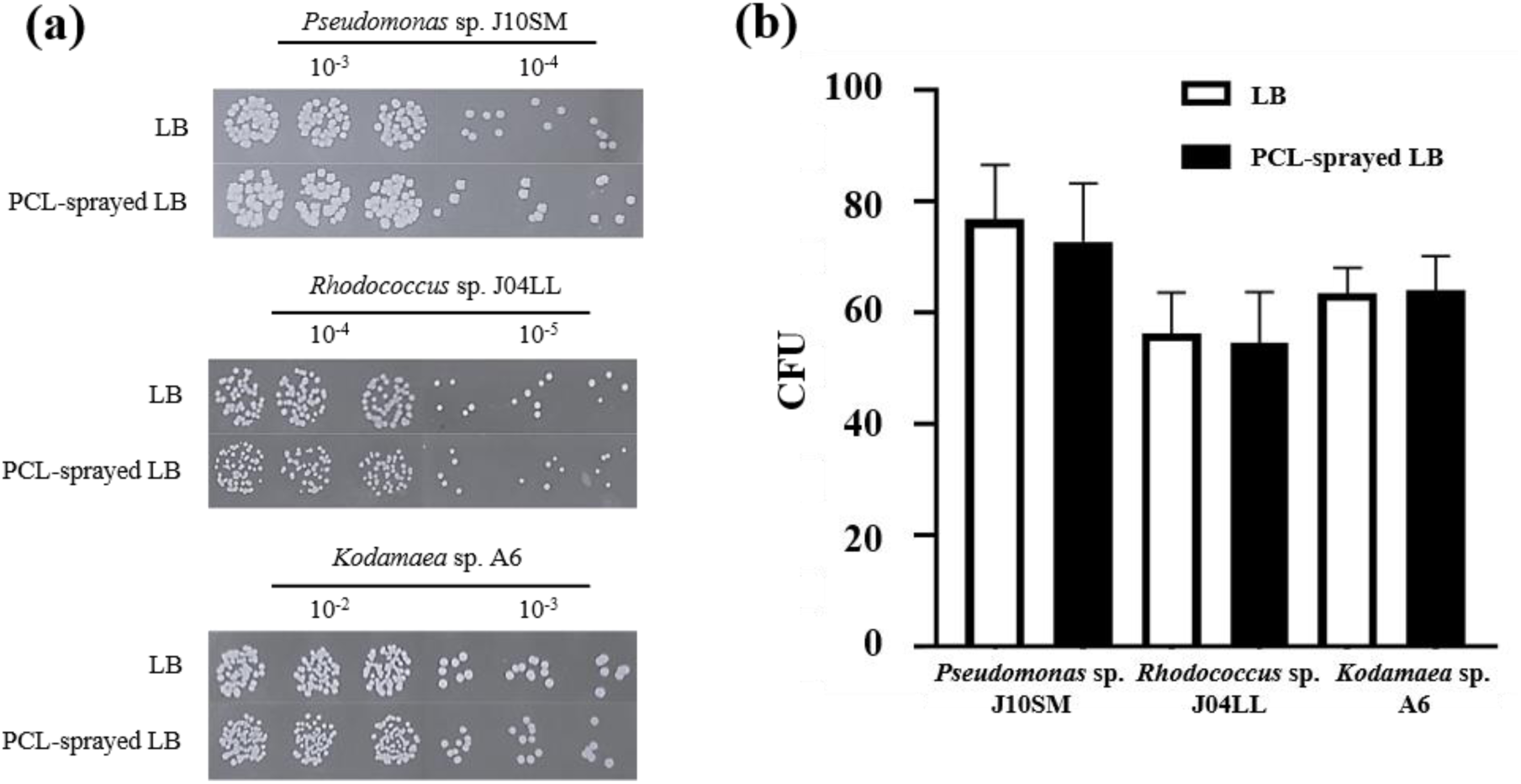
Toxicity assay of DCM-dissolved PCL sprayed on pre-inoculated plates. (a) Drop dilution assay for *Pseudomonas* sp. J10SM, *Rhodococcus* sp. J04LL and *Kodamaea* sp. A6 grown on LB agar and LB agar sprayed with PCL MPs (PCL-sprayed LB). (b) Bar chart showing the number of CFU counted on LB and PCL-sprayed LB for each strain (n=3). Data is presented as mean ± SD.

No apparent lethal effects were observed during the toxicity assays performed here on the selected strains. This lack of toxicity against bacterial and yeast cells is likely due to the inherent nature of the spraying method, where a rapid evaporation of the solvent occurs before plastic MPs reach the plate. In this regard, it has been suggested that the working distance plays a crucial role in obtaining solvent-free plastic particles. Therefore, it is probable that cells on pre-inoculated plates are not even exposed to DCM^43–45^ and, hence, popular inoculating techniques, such as spread plating or streaking, can be safely applied before spray-coating.

### Clear zone test: rapid screening of polyester depolymerization amongst microbial isolates

To showcase the spraying method’s value, a laboratory collection of 24 microbial isolates obtained from environmental samples (i.e., soil, freshwater reservoirs and compost) were screened for their ability to produce hydrolysis halos in all six polyesters (Table 2). Twenty-one out of the 24 isolates had been previously isolated as halo-forming microbes on BH solid media containing emulsified PCL (note: PCL is the only aliphatic polyester that allows an easy synthesis of emulsified solid plates). The screening provided 23 strains producing hydrolysis halos on PCL, eight on PES, eight on PBS, seven on PLA, six on PHB and two on PBAT (Table 2). Of all isolates, 58 % of them had the ability to hydrolyze more than one polyester, with PCL yielding -as expected from the origin of isolation- the highest number of degraders (96 %). The isolates, being mostly bacteria, were taxonomically classified and grouped into the *Actinomycetota*, *Bacillota* and *Pseudomonadota* phyla. These taxa contain genera that have commonly been reported for having plastic-degrading abilities.^12^ On the one hand, mesophilic isolates belonging to the *Streptomyces*, *Cellulomonas*, *Rhodococcus* and *Pseudomonas* genera are well-known for their resourceful enzymatic arsenal, which include numerous extracellular hydrolases such as cellulases, lipases or proteases, and for their promising bioremediation and plastic biodegradation potential.^46–49^ On the other hand, halo-forming thermophilic strains were more diverse and included species from the *Thermoactinomyces*, *Streptomyces*, *Saccharopolyspora*, *Thermobifida*, *Brevibacillus* and *Mycobacterium* genera. Some of these, *Thermobifida* and *Brevibacillus* have been reported as useful sources of enzymes for polyester biodegradation, like PCL or poly(ethylene terephthalate) (PET).^50,51^ The only fungus that demonstrated polyester-hydrolyzing activity was the basidiomycetes *Vishniacozyma* sp., which was able to grow optimally at 20 °C and hydrolyze all polyesters (including PBAT) at this temperature, with the exception of PHB.

**Table 2.** Collection of laboratory isolates with identified polyester-hydrolyzing capabilities.

|  | Isolate | Origin | Growth temperature (°C) | Polyesters<sup>a</sup> |  |  |  |  |  | Reference or source |
| --- | --- | --- | --- | --- | --- | --- | --- | --- | --- | --- |
|  |  |  |  | PBAT | PBS | PCL | PES | PHB | PLA |  |
| *Amycolatopsis orientalis* | DSM40040<sup>T</sup> | Soil | 28-37 |  |  |  |  | + | + | Ref.<sup>52</sup> |
| *Brevibacillus* sp. | SC8 | Compost | 50 |  | + | + |  |  |  | This study |
| *Cellulomonas* sp. | G3C1b | Freshwater pond | 30 |  | + | + | + |  |  | This study |
| *Cellulomonas* sp. | G4A2 | Freshwater pond | 30 |  | + | + | + |  |  | This study |
| *Mycobacterium* sp. | SF82 | Compost | 50 |  |  | + |  |  |  | This study |
| *Rhodococcus* sp. | J04LL | Soil | 30 |  | + | + | + |  | + | This study |
| *Saccharopolyspora* sp. | E50 | Soil | 50 |  |  | + |  |  |  | This study |
| *Streptomyces* sp. | J04ES | Soil | 30 |  |  | + |  | + |  | This study |
| *Streptomyces* sp. | J06LL | Soil | 30 |  |  | + |  |  |  | This study |
| *Streptomyces* sp. | AC1 | Soil | 50 |  |  | + |  |  |  | This study |
| *Streptomyces* sp. | C3a | Soil | 50 |  |  | + |  |  |  | This study |
| *Streptomyces* sp. | A3D2 | Soil | 30 |  |  | + |  | + |  | This study |
| *Streptomyces* sp. | A6a1 | Soil | 30 |  |  | + |  | + | + | This study |
| *Streptomyces* sp. | A6b | Soil | 30 |  |  | + |  | + | + | This study |
| *Streptomyces* sp. | J01BES | Soil | 30 |  |  | + |  |  |  | This study |
| *Streptomyces* sp. | J02LL | Soil | 30 |  |  | + |  |  | + | This study |
| *Streptomyces* sp. | J20SM | Soil | 30 |  |  | + |  |  |  | This study |
| *Streptomyces* sp. | J21SM | Soil | 30 |  |  | + |  |  |  | This study |
| *Stutzerimonas* sp. | P7301 | Soil | 30 |  | + | + | + |  | + | This study |
| *Stutzerimonas* sp. | P730a | Soil | 30 |  | + | + | + |  |  | This study |
| *Thermoactinomyces* sp. | A3a | Soil | 50 |  |  | + |  |  |  | This study |
| *Thermobifida fusca* | DSM 43793 | Mushroom compost | 50 | + |  | + | + | + |  | Ref.<sup>53</sup> |
| *Thermobifida* sp. | P150a | Soil | 50 |  | + | + | + |  |  | This study |
| *Vishniacozyma* sp. | Blapla | Airborne | 20 | + | + | + | + |  | + | This study |

<sup>a</sup> Symbol + indicates hydrolysis halos formed by each microbial isolate in the corresponding polyester-sprayed plates.

Hydrolysis halos were observed for all polyesters (Figure 3) after 2-5 days of incubation, although two weeks were required for the formation of hydrolysis halos in PBAT by *Vishniacozyma* sp. at 20 °C. Considering that PBAT is an aromatic-aliphatic polyester, it is probable that the presence of an aromatic ring in its chemical structure could create a steric interference that prevented hydrolases from easily attacking the polymer ester bonds and thus reduce the hydrolysis rate.^54^ Our positive control for PBAT degradation *Thermobifida fusca* DSM 43793^53^ was able to hydrolyze this polyester after only 2 days of incubation at 50 °C. This difference in hydrolyzing capabilities between strains might be a result of each strain’s intrinsic enzymatic potential to hydrolyze PBAT or as a consequence of the incubation temperatures (Table 2). The activation energy of an enzymatic reaction is highly dependent on temperature, with higher temperatures reducing the activation energy threshold and favoring higher reaction rates.^55^ Additionally, in the case of plastics, intermolecular forces between polymer chains are weakened when the material is heated and the polymer bonds become more accessible to enzymatic attack.^56^ Therefore, it is logical that PBAT hydrolysis is enhanced at higher temperatures and thermophilic strains are in a more advantageous position to hydrolyze plastics.

**Figure 3.**
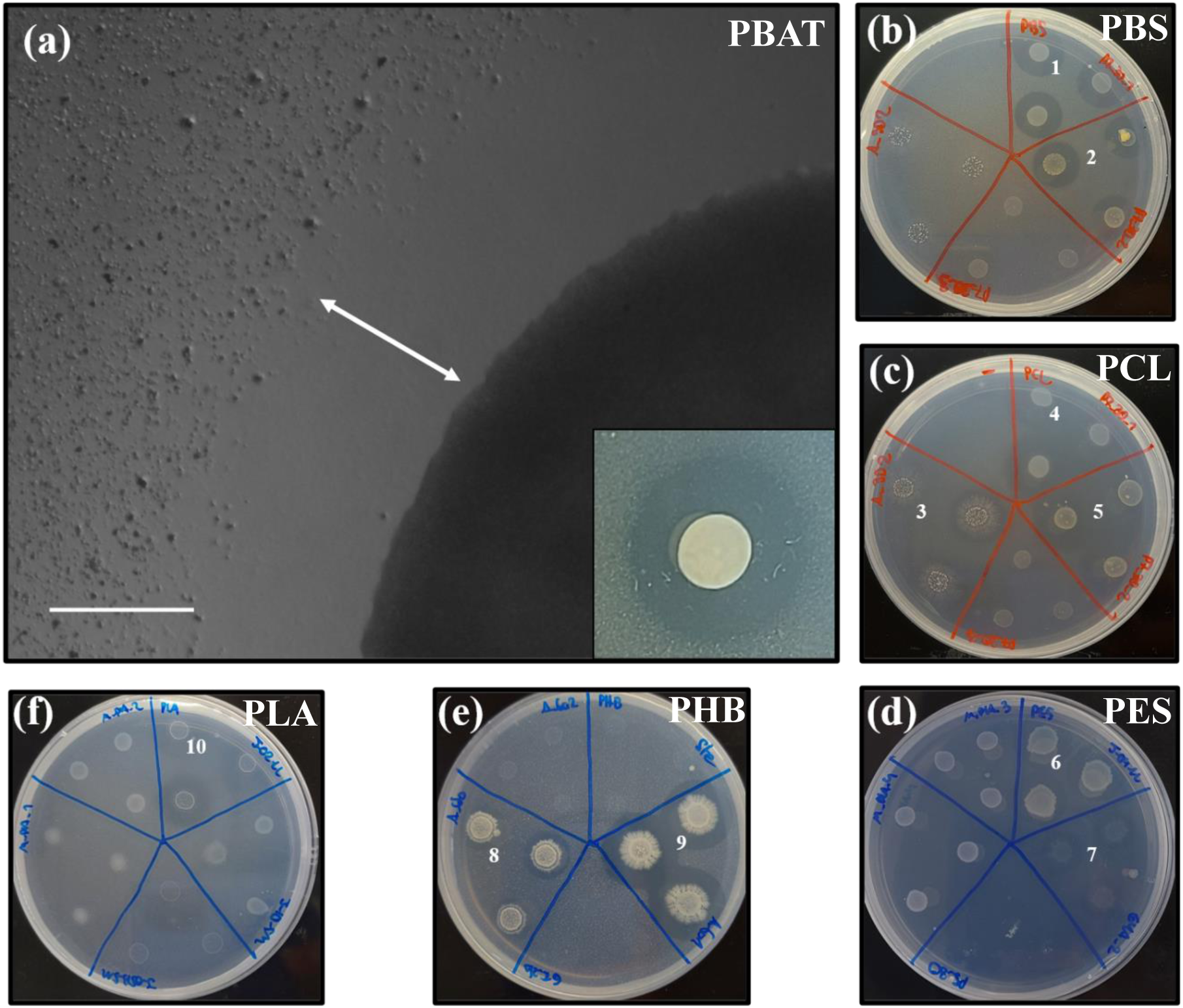
Polyester clear zone tests with different isolates. (a) Light microscope image of a PBAT- hydrolyzing colony corresponding to isolate Blapla. An image of the same halo observed by plain sight can be observed at the bottom right corner. The white arrow indicates the degradation halo width. Scale bar: 100 µm; (b) PBS-hydrolyzing strains: P7301 (1) and P730a (2); (c) PCL- hydrolyzing strains: A_3D2 (3), P7301 (4) and P730a (5); (d) PES-hydrolyzing strains; J04LL (6) and G4A_2 (7); (e) PHB-hydrolyzing strains: A6b (8) and A3D2 (9); (f) PLA-hydrolyzing strains: J02LL (10).

The hydrolysis of the MPs at the colony edges was visualized under a microscope (Figure 3a). Interestingly, an initial loss of opacity and shape of the MPs was observed during the first days of incubation (Figure S4). This was attributed to an initial increase in porosity as a result of enzymatic hydrolysis of the amorphous regions of the plastic. As pores grow, water flows freely into the particles and, consequently, the refractive index is diminished, increasing transparency.^57^ In addition, MP hydrolysis was not restricted to the region outside the colony boundaries, as particles within the inoculation spots were also degraded, facilitating the detection of halo-forming isolates during early stages of incubation. Such evaluation was also useful for strains that grow masking the hydrolysis halo, such as fungi or members of the phylum *Actinomycetota* like *Streptomyces* spp. Some of these microorganisms produced hydrolysis halos that were more difficult to detect by sight as these were covered by the growth of the strain itself (Figure 3c).

### Screening for PCL-hydrolyzing isolates in compost: a comparison of methods

The spraying method was tested for screening new PCL-degrading microbes in an environmental sample, i.e. homemade compost soil, and compared with plates generated with emulsified PCL. The main objective was to evaluate how polyester availability -either emulsified in the medium or in the form of polyester MPs on the agarose surface-affected the screening sensitivity of each method. PCL was selected because of *i*) its elevated biodegradability by environmental microorganisms, and *ii*) its high solubility in acetone, being one of the few polyesters that is easily emulsified in solid medium without the use of an ultrasonic processor and the addition of undesired surfactants.^58^

Colony counting revealed that, after three weeks, a similar number of colonies had grown on both media (PCL-sprayed medium: 168.2 ± 28.7 CFU; PCL-emulsified medium: 145.7 ± 16.4 CFU; Student’s t-test; p = 0.1258), illustrating again how the spraying method does not prompt cell death (Figure 4). Nevertheless, the number of halo-forming colonies was significantly lower in the case of PCL-sprayed plates (2.5-7.9 % of all colonies were able to hydrolyze sprayed PCL MPs) *vs* PCL-emulsified plates (10.2-17.6 % of all colonies produced hydrolysis halos; Student’s t-test; p < 0.002).

**Figure 4.**
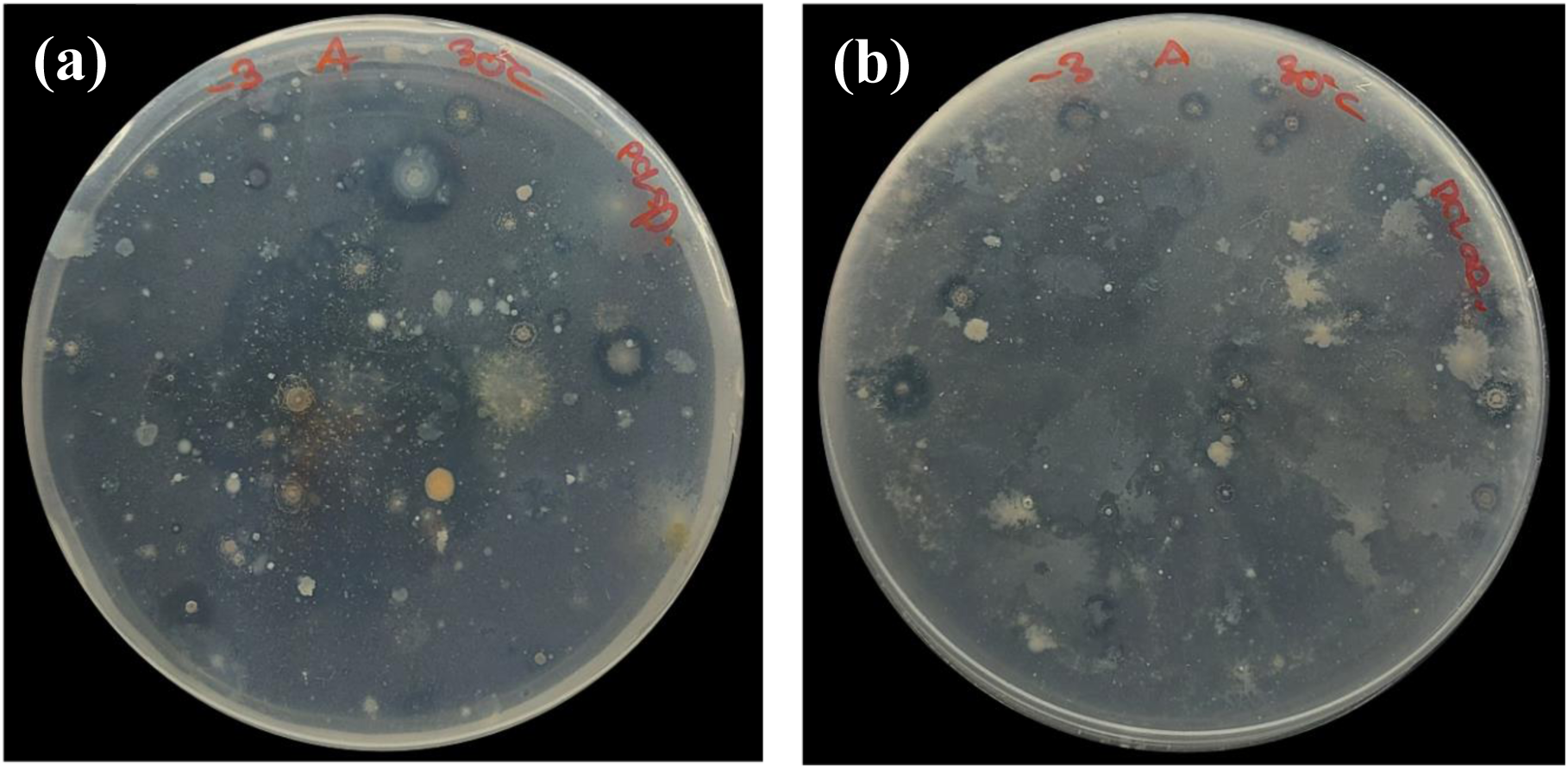
PCL hydrolysis halos formed by microbial colonies from compost. (a) PCL-sprayed solid BH and (b) solid BH containing 0.1 % w/v emulsified PCL are compared.

While the spraying method generated MPs of approximately 14 µm in diameter, those within emulsified polyesters can be under 1 µm, depending on the preparation method.^59^ Here, particles formed by PCL-acetone emulsions had an approximate median size of 200 nm according to SLS measurements (Figure S5). Particle size plays a vital role during polymer biodegradation as extracellular hydrolases are unable to penetrate the polymer matrix and their cleavage activity is restricted to the particle’s surface.^54^ Consequently, the specific surface area (SSA) -defined as the total surface area per unit mass-is essential in controlling the adhesion and hydrolysis kinetics during polyester depolymerization. Hence, in enzymatic assays and biodegradation tests, smaller plastic fragments are more rapidly degraded. This was shown in studies conducted on plastic particles in the nano- and micrometric scale range, as well as larger fragments, such as entire pellets or films.^60–63^ In addition, other polymer properties, such as hydrophobicity, might play a more relevant role in the depolymerization of larger particles. In this case, a more hydrophobic plastic would hinder the adsorption of water, essential during hydrolytic reactions.^59^

The estimated SSA value for PCL emulsions prepared here (13.09 m^2^/g) was 35 times higher than that of PCL-sprayed preparations (0.37 m^2^/g) (Table S2). The large particles generated during the spraying method (i.e. 14 µm) and the resulting lower SSA might be a limiting factor hampering depolymerizing rates. Furthermore, the actual SSA available for enzymatic hydrolysis might be reduced even more, as sprayed MPs are spread over the surface of the medium rather than fully engulfed in the agarose matrix like emulsified PCL particles. Under these conditions, enzymatic hydrolysis of the MPs will become more reliant on some factors such as water and enzyme diffusion into the particles. Nevertheless, the *a priori* reduced sensitivity of the spraying technique may offer a stricter screening platform for more efficient polyester degrading microbes, possibly with much higher depolymerizing potential. The discrimination amongst isolates with different degrading capabilities could offer a new perspective for understanding plastic degradation dynamics in the environment.

### Polyester-hydrolyzing potential across multiple environments

The simplicity of our plate spray-coating method allowed a broad functional assessment of the microbial degrading potential for all six polyesters across 15 different locations that included agricultural soils, compost, as well as freshwater and marine sediments (Figure 5). The hydrolyzing potential of protein (i.e. casein from milk) was used as a natural labile polymer control. This comprehensive screening established a clear ranking of polymer biodegradability across all environments, from more to less degradable: protein, PHB, PCL, PES, PBS, PLA and PBAT. Most interestingly, it revealed a notable lack of plastic biodegrading potential in marine environments, being up to three orders of magnitude lower than in terrestrial systems (Figure 5).

**Figure 5.**
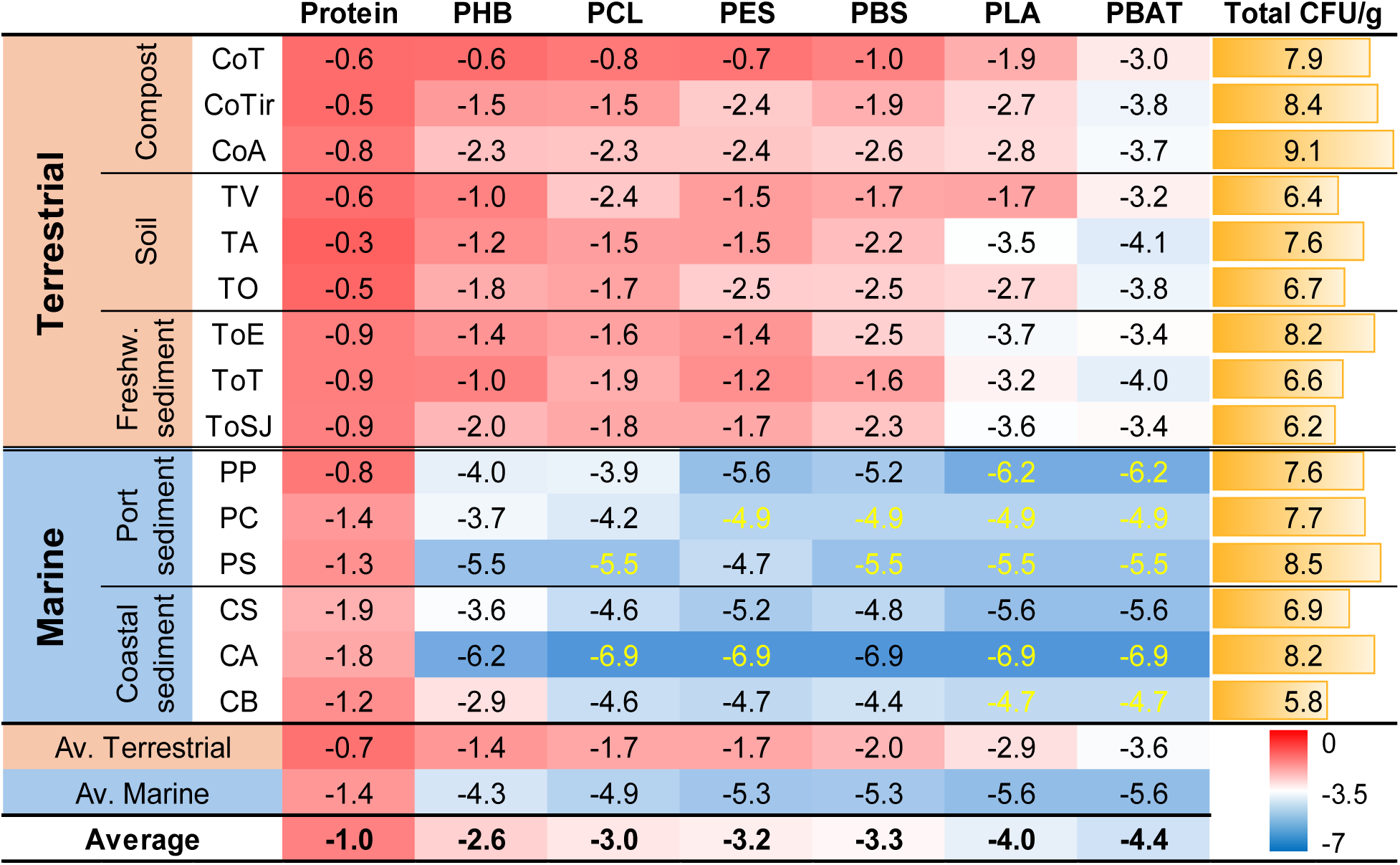
Hydrolysis halo frequencies for each polymer across different environments. Frequencies were calculated by dividing the number of halo-forming colonies by the total culturable microbes grown with labile substrates. Frequency values and total CFU/g are expressed in logarithmic scale (log_10_). Numbers highlighted in yellow represent values below the detection limit (no halos were detected).

As expected, protein hydrolyzing microbes abounded across all samples, averaging 10.9% of all plate-growing microbes (Figure 5). Exoproteases have been reported as one of the most common extracellular enzymes in a great variety of natural settings, especially in environments rich in organic matter.^64–66^ A much lower frequency of plastic degraders was obtained, although this varied greatly between polyesters, i.e. ranging from 10^-2.6^ for PHB to below 10^-4.4^ for PBAT (Figure 5). This meaning that, amongst culturable bacteria, there was one in every nine capable of protein hydrolysis, one in 378 of PHB hydrolysis and one in over 26,000 of PBAT hydrolysis. PHB showed the highest biodegradability (i.e. 10^-2.6^) and was the only polyester for which hydrolysis halos were observed in all environmental samples. This is not surprising considering that PHB is an aliphatic polyester that serves as a natural carbon storage for many bacteria in different environments and, hence, secreted extracellular PHB depolymerases abound in nature.^67^ PHB materials have shown remarkable biodegradation in a wide range of natural settings, including terrestrial and marine ecosystems.^68,69^

Despite being of synthetic origin, PCL, PES and PBS also showed a high frequency of microbial degraders across the different environments tested (i.e. frequencies averaging 10^-3.0^, 10^-3.2^ and 10^-3.3^, respectively; or one of every 1003, 1428 and 2154 culturable microbes were capable of PCL, PES and PBS hydrolysis, respectively). A crucial trait shared by these polyesters is their low glass transition temperature (T_g_) (PCL, −60 °C; PES, ca. −10 °C; PBS, −20 °C), which promotes greater chain flexibility and water diffusion into the polymers under mesophilic conditions.^54^ As a result, enzymatic catalysis is facilitated, leading to higher depolymerization rates. This, together with the considerable frequency of hydrolyzation observed for all three polyesters in our study would explain why these plastics are biodegradable under most natural conditions, with marine environments being the least favorable as will be discussed below.^69–73^ Given the structural similarities between PCL, PES and PBS, and the substrate promiscuity associated with esterases, it is tempting to suggest that one same enzyme could hydrolyze all three polymers.^74^ However, some polyesterases are highly sensitive to the carbon chain length of the monomeric components surrounding the ester linkage.^75,76^ Furthermore, beyond each depolymerase’s own substrate affinity, polyester hydrolysis is also controlled by the physical properties of each polymer, such as crystallinity and molecular weight.^77–79^

PLA and PBAT were the least biodegradable materials of those tested in this study, i.e. with frequencies of hydrolyzing microbes below 10^-4.0^ and 10^-4.4^, respectively (meaning less than one of every 9300 and 26,000 culturable microbes could hydrolyze PLA and PBAT, respectively; Figure 5). Most disturbingly, while PLA- and PBAT-degrading microbes could be detected in terrestrial and freshwater environments, these remained below detection limits in all marine samples (under 10^-4.3^ - 10^-6.9^). Low PLA degrading potential in the environment has been previously reported.^17,80^ Presumably, PLA’s high hydrophobicity and Tg (60 °C) make this polyester difficult to biodegrade in the environment beyond industrial composting.^81^ PLA hydrolysis is usually conducted by two types of depolymerases: lipases and proteases, the latter showing a high selectivity for PLA and no other polyesters.^82^ More importantly, a high degree of enantioselectivity against poly(L-lactic acid) or PLLA was observed. This phenomenon might be the result of the similarity between the monomeric component of PLLA, i.e. L-lactate, and L-alanine in proteins.^83^ The frequency of exoprotease producers calculated in this work was 10^3^ times higher than that of PLA-hydrolyzing microbes, hence, inferring that exoprotease production clearly does not imply PLA hydrolysis under mesophilic conditions.

PBAT was the only aliphatic-aromatic polyester used in this work and the material that showed the lowest biodegrading potential amongst the environments explored (Figure 5). PBAT is a linear random copolymer consisting of: *i*) an aliphatic section composed of 1,4-butanediol and adipic acid (BA) that is recognized as the flexible or soft unit of the polymer, and *ii*) a rigid aromatic section made of 1,4-butanediol and terephthalate units (BT) that is the main element of the polymer’s crystalline region.^84,85^ It is known that the presence of aromatic regions in a polymer structure may pose a steric hindrance for enzymatic attack, beyond reducing the flexibility of the polymer chain and restricting the accessibility to the ester bond.^86^ For such polymers, a concentration of the aromatic component (i.e., terephthalic acid) below 50 mol% is fundamental for biodegradation to happen.^87^ On the other hand, the BA region, due to its flexibility and contribution to lowering the material’s Tg, is regarded as the main component of the amorphous region which is subject to enzymatic hydrolysis.^57^ However, under mesophilic conditions, the BT crystallinity outweighs the contribution of the BA region to the final biodegradability of the polymer. Such constraints are overcome at higher temperatures, such as under industrial thermophilic composting conditions, when polymer chains move more freely and ester bond hydrolysis of the BT region is enhanced.^88^ This would explain why thermophilic PBAT degraders are more common and therefore more frequently isolated, although PBAT depolymerization at lower temperatures has also been reported.^89–91^ Since PBAT degradation is clearly hampered under low temperatures, finding and characterizing efficient mesophilic PBAT hydrolases is of high biotechnological interest.^75,92,93^

Marine environments presented around 1,000 times lower polyester degrading potential than terrestrial ones, being in many cases even below the method’s detection limit (Figure 5). Interestingly, this was not the case for the protein control, where marine degrading potential was only 5.4 times lower, suggesting ecological and not methodological factors involved. The low plastic biodegrading potential in marine ecosystems has been previously suggested but never shown so clearly, systematically and functionally as in the present study.^72,94,95^ Biodegradation rates in marine environments are generally regarded as suboptimal as a result of lower temperatures, reduced oxygen levels and higher salinity as well as reduced microorganism concentrations.^9,96^ However, while most of these hampering factors may hold true for microbial communities living in the water column in colder latitudes, having analyzed sediments from Mediterranean ports and coastal beaches rules out factors such as temperature (seawater temperatures of 20-22 °C, compared to freshwater sediments i.e. 16-20 °C) and microbial density (see Figure 5). Such deficiency of polyester degrading potential in marine environments is likely due to a combination of factors, such as: *i*) a lack of esterolytic activity (i.e. carboxylesterases, EC 3.1.1.1; lipases, EC 3.1.1.3; and cutinases EC 3.1.1.74) as a consequence of the lower exposure to complex plant-derived detritus in marine environments; *ii*) the high ionic force of seawater, hampering the interactions between hydrophobic compounds, such as plastics, and the enzyme’s active sites; or *iii*) distinct ecological strategies selected in marine environments, e.g. a possible predominance of surface-anchored hydrolases -ectoenzymes-that promote ‘selfish’ degrading strategies and that would not be detected with the “clear zone” method employed here.^97–99^ In addition, it is important to keep in mind that all experiments were carried out under aerobic conditions and, therefore, the role of anaerobic microorganisms —present in sediments— in polyester degradation was not considered.

## CONCLUSIONS

The low-cost and user-friendly airbrush technique for dispersing polyester microplastics (MPs) on solid media enabled an extensive screening of the biodegradation potential of various polyesters across diverse environments. The expected ranking of polyester biodegradability was confirmed (i.e., PHB > PCL > PES > PBS > PLA > PBAT). However, the study underscored a striking disparity in biodegradation potential between environments, with marine ecosystems showing degradation rates that were three orders of magnitude lower than their terrestrial counterparts. This significant gap, coupled with the fact that marine environments are major sinks for plastic pollutants, raises concerns about the limited abundance of plastic-degrading organisms in marine ecosystems. Our accessible method opens up new possibilities for future ecological research and facilitates the rapid isolation of plastic-degrading microbes with promising biotechnological applications.

## ASSOCIATED CONTENT

The plate spraying protocol; spraying equipment and recommendations; further analysis of sprayed plates; sprayed and emulsified particles different size, polyester hydrolysis halos and frequencies and equations for the calculation of SSA are available as supplementary material free of charge.

## AUTHOR INFORMATION

### Corresponding Authors

**Alberto Contreras Moll** - *Department of Biology, University of the Balearic Islands, Palma 07122, Spain*;

**Joseph Alexander Christie-Oleza** - *Department of Biology, University of the Balearic Islands, Palma 07122, Spain*;

### Author Contributions

A.C.-M. planned and performed the experiments, analyzed the data and wrote the manuscript. T.O.-V. contributed to the conceptualization and revision of the paper. RDI.M and M.A.-F. assisted in the execution of the experiments. B.N. and R.B. aided in revision of the paper and supervision of the project. J.A.C.-O. was involved in funding acquisition, project management, interpretation of the results, and preparation of the manuscript.

The manuscript was written through contributions of all authors. All authors have given approval to the final version of the manuscript.

### Funding Sources

This work was supported by the research projects polyDEmar (PID2019-109509RB-I00 and PDC2022-133849-I00), AlivePlastics (TED2021-129739B-I00) and plasticROS (PID2022-139042NB-I00) funded by MICIU/AEI/10.13039/501100011033 and EU NextGenerationEU/PRTR. A.C.-M. was supported by the FPU21/05173, and T.O.-V. by FPU19/05364 contract from the Spanish Ministry of Universities. J.A.C-O. was supported by the Ramón y Cajal contract RYC-2017-22452 (funded by MCIN/AEI/10.13039/501100011033 and “ESF Investing in your future”).

## ACKNOWLEDGMENT

The authors would like to thank the students from high schools I.E.S Josep Maria Llompart, I.E.S. Josep Font i Trias and I.E.S Santa Maria, as well as the program AMGEN TransferCiencia and the FCRI for organizing the workshops as a proof of concept. Authors acknowledge TIRME, S.A. for providing the industrial compost used in this study. The authors also thank Dr. Antonio Busquets Bisbal for the acquisition and analysis of the SEM and HRSTEM images and Dr. José F. González Morey for the technical support and guidance provided during the SLS analysis of the emulsified PCL particles.

## ABBREVIATIONS

BA: 1,4-butanediol-adipate
BH: Bushnell-Haas
BT: 1,4-butanediol-terephthalate
CFU: colony forming units
DCM: dichloromethane
HRSTEM: high-resolution transmission electron microscopy
LB: Luria-Bertani
MP: microparticle
PBAT: poly(butylene adipate-co-terephthalate)
PBS: poly(butylene succinate)
PCL: polycaprolactone
PCR: polymerase chain reaction
PES: poly(ethylene succinate)
PET: poly(ethylene terephthalate)
PHB: poly[(R)-3-hydroxybutyrate]
PLA: poly(lactic acid)
rDNA: ribosomal deoxyribonucleic acid
SEM: scanning electron microscopy
SLS: static laser light scattering
Tg: glass transition temperature.

## Notes

### Competing Interest Statement

The authors have declared no competing interest.

