## Supplementary Information for "Lack of functional polyester-biodegrading potential in marine *versus* terrestrial environments evidenced by an innovative airbrushing technique"

### Table of Contents

**Plate spraying protocol** **S3**

**Figure S1.** Spraying equipment and recommendations **S10**

**Figure S2**. Plate opacity of different polyester-sprayed plates **S11**

**Figure S3**. Violin plot distribution of polyester microparticles according to their diameter **S12**

**Figure S4**. PBAT hydrolysis halo around a small colony of *Vishniacozyma* sp. **S13**

**Figure S5**. Median size of PCL particles obtained from an emulsified solution **S14**

**Table S1**. Descriptive statistical values associated with all polyester microparticle samples **S15**

**Table S2**. Equations and data used to calculate the estimated SSA values for the air-spraying and emulsification methods. **S16**

**Plate spraying protocol: screening polyester-degrading microorganisms**

This protocol describes the use of an artist's airbrush as a simplified spraying equipment to prepare polyester-containing solid media. To this end, the *Preparation of polyester dissolutions* alongside the following *Plate spraying* are included. Additionally, the protocol comprises a final section (*Airbrush equipment cleaning*) dedicated to the airbrush maintenance and envisioned to allow users to make the most of the equipment.

*Preparation of polyester dissolutions*

Due to their high hydrophobicity, plastics are normally effectively dissolved in organic solvents, such as acetone or chloroform. In this work, dichloromethane was selected as the appropriate organic solvent due to its high volatility, allowing a homogeneous dispersion of polyester microparticles while spraying, as well as limiting the exposure of the microorganisms to the solvent. Furthermore, the specific physicochemical properties of the polymer used will vary between brands (e.g., molecular weight, crystallinity, etc.) and forms (e.g., granulated, powder, film, etc.) that will inevitably affect the dissolution process. As a result, additional steps like controlled heating or overnight stirring to ensure the plastic dissolution could be implemented. However, this particular protocol relies solely on external sonication and mechanical stirring to minimise the dissolution time and avoid unnecessary risks related to solvent heating.

**CAUTION**: highly volatile solvents must be handled in a fume hood at all times to prevent vapour inhalation, especially with potentially carcinogenic substances such as dichloromethane (DCM). This is also necessary when following the steps detailed in the *Plate spraying* section. In addition, glassware is recommended over plastic equipment when working with chlorinated organic solvents.

*Materials*

- Methylene chloride/dichloromethane 99 % GLR stabilised with amylene (Labkem or equivalent)

- Polyesters:
    - Poly(ε-caprolactone) or PCL flakes ($M_w$ ~14k, $M_n$ ~ 10k) (Merck KGaA or equivalent)
    - Ingeo 2003D grade polylactic acid or PLA granules ($M_w$ ~ 180k, $M_n$ ~ 100k) (~ 4.3 % D-lactate) (NatureWorks LLC or equivalent)
    - Polyethylene succinate or PES fragments ($M_w$ ~ 10k, $M_n$ ~ n/s) (Merck KGaA or equivalent)
    - Poly[(R)-3-hydroxybutyric acid] or PHB powder ($M_w$ ~ n/s, $M_n$ ~ n/s) (Merck KGaA or equivalent)
    - Polybutylene succinate or PBS granules extended with 1,6-diisocyanatohexane ($M_w$ ~ n/s, $M_n$ ~ n/s) (Merck KGaA or equivalent)
    - Poly(butylene adipate-co-terephthalate) or PBAT pellets ($M_w$ ~ 130k, $M_n$ ~ 49k) (EcoWorld® PBAT 2208) (JinhuiZhaolong Co., Ltd or equivalent)
- Glassware (borosilicate glass bottles, glass pipettes, beaker)
- Chemical-resistant gloves for protection against organochlorine solvents (e.g., polyvinyl alcohol or PVA gloves) (ALPHATEC® 15-554 from Ansell™ or equivalent)
- Ultrasonic bath

1- Prepare a 1.5 % w/v dissolution by adding 0.3 g of polyester pellets/flakes/powder to 20 mL of DCM in a glass bottle.

*More concentrated solutions (i.e., 3.0 % w/v) can be prepared for easily soluble polyesters like PCL or PES to increase plate opacity, which is essential to identify halo-forming colonies, whilst maintaining the spraying volume (1 mL).*

2- Seal all glass bottles with a secure cap and label them accordingly.

*Regarding DCM, it is catalogued as a health hazard (GHS 8) due to its suspected carcinogenicity and as a harmful substance (GHS 7) prone to cause irritation when in contact with the skin or eyes, as well as dizziness when inhaled.*

3- Optional step: subject all dissolutions to external sonication in an ultrasonic bath at 50/60 Hz for at least 15 minutes to accelerate polyester dissolution.

*Sonication time will vary depending on the polyester type, form and intrinsic solubility in the solvent (i.e., PBS and PLA: 30-40 min.; PBAT, PCL, PES, PHB: 10-20 min.). Furthermore, in the case of PHB, it was found that a preceding step consisting in mechanical stirring for 30 minutes improved powder disaggregation and its dissolution in the following sonication step.*

4- Do not open glass bottles immediately after sonication. Let them stand for at least 15 minutes before being used.

*An increase of the liquid temperature inside the bottle is expected to take place when subjecting it to sonication in an ultrasonic bath. Considering that the liquid, DCM, is a solvent with a low boiling point, the pressure inside the container is expected to increase. This is clearly evident for dissolutions that have been sonicated for over 30 minutes, like PLA or PBS dissolutions. Therefore, it is crucial to leave enough headspace inside glass containers (i.e., only 20 mL of solution are prepared in 100 mL glass bottles) and not to open them immediately after sonication.*

5- Store the solution in a dry, cold, well-ventilated place away from direct sunlight and heat or ignition sources until needed.

*Plate spraying*

Inoculation by plate streaking, plate spreading or drop deposition can be done previous to the spraying step as the exposure to the solvent does not seemingly affect cell viability (see *Assessment of solvent toxicity on inoculum viability during plate spraying applications* in the Results and Discussion section).

*Materials*

- Airbrush pistol (e.g., double action gravity-fed FE-130 airbrush, Fengda® or equivalent)

  *The airbrush model used in this work is simply a recommendation and can be replaced by any other suitable airbrush model.*

- Air compressor (e.g., FD18-2K air compressor, Fengda® or equivalent)

- Transparent carbon-free solid medium (e.g., Bushnell-Hass medium or other carbon-free basal medium with agarose).

- Petri plates

1- Assemble the spraying equipment and connect the air hose to the air compressor and the airbrush gun.

*Sealing the air connections between the airbrush gun, the air hose and the air compressor with polytetrafluoroethylene (PTFE) tape is highly recommended in order to avoid a pressure loss due to air leaks.*

2- Turn it on and wait for the pressure to build up to the maximum value (set at 4 bar by default).

*The FD18-2K air compressor includes an auto-stop/start function that controls the air intake and activates whenever the pressure descends to 3 bar (by default) when spraying. This automatic function keeps the pressure (i.e., working pressure) stable during spraying sessions and raises it back to its maximum value when the airbrush is not being used.*

3- Make sure the airbrush works properly by spraying some air.

*Additionally, a few millilitres of water can be sprayed through the nozzle to check for bubbling or sizzling as air leakage indicators.*

4- Place the airbrush gun and the rest of the material needed inside the fume hood: glassware, polyester dissolutions, inoculated Petri plates and the DCM container.

5- Carry out an initial cleaning by pipetting 3-4 mL of DCM into the airbrush paint/fluid cup using a glass pipette.

6- Spray the solvent inside a glass beaker to clean the inside of the airbrush head and blow out any remaining polyester traces from previous spraying rounds.

7- Add 1 mL of the desired polyester dissolution into the airbrush fluid cup by pipetting.

8- Uncover the Petri dish and spray the polyester dissolution in a circular manner onto the solid media. Cover the plate to avoid contamination. Make sure to spray the whole volume since some residual droplets might remain in the paint cup after the initial spraying.

*Keeping an adequate distance between the plate and the nozzle while spraying is essential to ensure that the solvent does not reach the surface of the plate. Thus, a distance of 15 cm is recommended for the polyesters and the airbrush system used in this work. Spraying time is usually between 3-4 seconds for 1 mL of DCM at a working pressure of 3 bars.*

*Additionally, when using a double action airbrush, to avoid the formation of residual droplets onto the nozzle tip, it is important to first slide back the trigger into its initial position -to stop the liquid release- before releasing it to stop the airstream.*

9- Place the closed Petri dish aside and let any traces of DCM evaporate in the fume hood for at least 30 minutes before placing them for incubation.

10- Before turning off the air compressor, spray ~3-4 mL of DCM into an empty beaker to clean out the airbrush inner channels.

*If more than one polyester dissolution is going to be blown through the airbrush pistol consecutively, it is recommended that a few millilitres (~3-4 mL) of DCM are sprayed in between spraying rounds to clean any remaining traces of previous polymers.*

11- Keep blowing out air for a few seconds after the entire solvent volume has been sprayed to minimise the amount of residual DCM inside the airbrush.

12- Set the airbrush aside, switch off the air compressor and let out the remaining pressurised air inside the hose by 1) blowing it out through the airbrush tip until pressure gradually descends to zero bars or 2) by emptying the moisture trap.

*A moisture trap is a useful accessory that captures water from the incoming air and helps reduce its total humidity rate. It is especially handy in humid environments to prevent water from accumulating inside the air hose and eventually being sprayed out through the nozzle.*

13- Clean the equipment by following the instructions given in the *Airbrush equipment: cleaning* section (see below)

*Airbrush equipment cleaning*

An adequate equipment maintenance is essential to extend the airbrush lifespan, which can be severely reduced by the constant use of organic solvents and an inadequate cleaning.

*Cleaning*

Keeping a regular cleaning routine is crucial to remove any remaining traces of organic solvents inside the airbrush that, otherwise, would slowly damage the internal rubber components, such as the O-rings.

*Materials*

- Cotton pads and buds
- Distilled water
- Detergent
- Airbrush cleaning brushes

*Basic cleaning*

1- Mix some regular dish washer detergent and distilled water to prepare a cleaning solution.

2- Fill the paint cup to the brim with the detergent solution and spray it. Repeat a second time.

3- Repeat the previous step a few times using only distilled water to rinse the airbrush paint cup and passages.

4- Ensure all water has been blown out by spraying air for a few more seconds.

5- Disassemble the airbrush and let any water remnants dry out.

*Thorough cleaning*

1- Perform a basic cleaning.

2- Disassemble the airbrush.

3- Dip a cotton pad or a cotton bud in the cleaning solution.

4- Use cotton pads to gently clean the needle and wash away any plastic debris found on the outside of the airbrush.

5- Cotton buds and soft brushes are useful to reach and successfully clean other areas such as the inside of the paint cup or the needle passage.

*Additionally, when encountering dried plastic patches that are difficult to wash away, it is recommended using soft brushes such as the designated airbrush cleaning brushes to scrape them off.*

6- The interior of the nozzle can be cleaned up by wetting the needle tip in the detergent solution and gently inserting it in the nozzle and turning. Alternatively, a piece of paper rolled into a point or a sewing needle can be used to complete this task.

7- Dip the airbrush components in a water-filled container to rinse them. Then, make sure to dry them thoroughly.

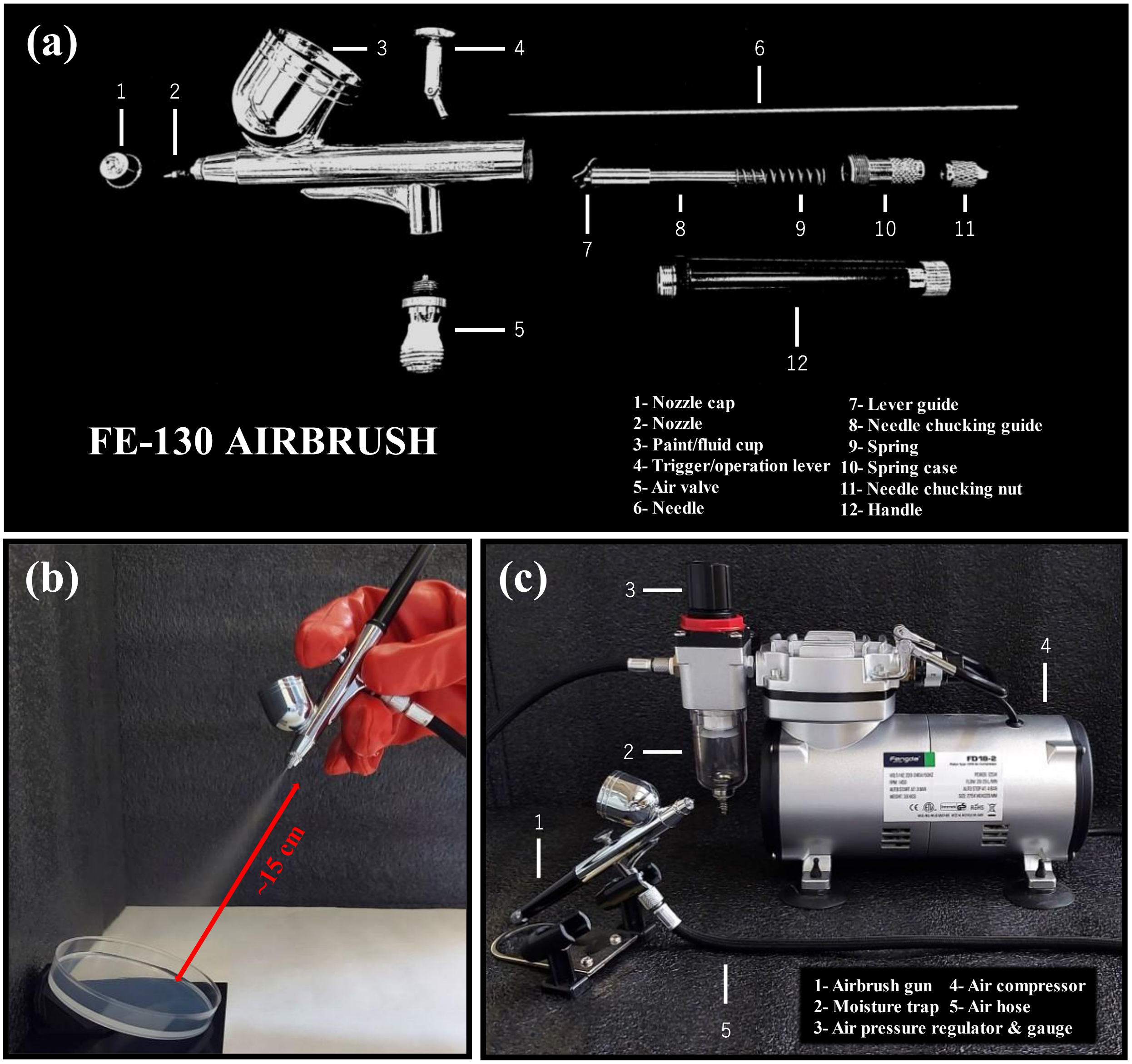


**Figure S1**. Spraying equipment and recommendations. (a) FE-130 airbrush parts diagram. (b) Plate spraying distance between the agar plate and the nozzle. (c) Double action airbrush gun, air compressor and additional components.

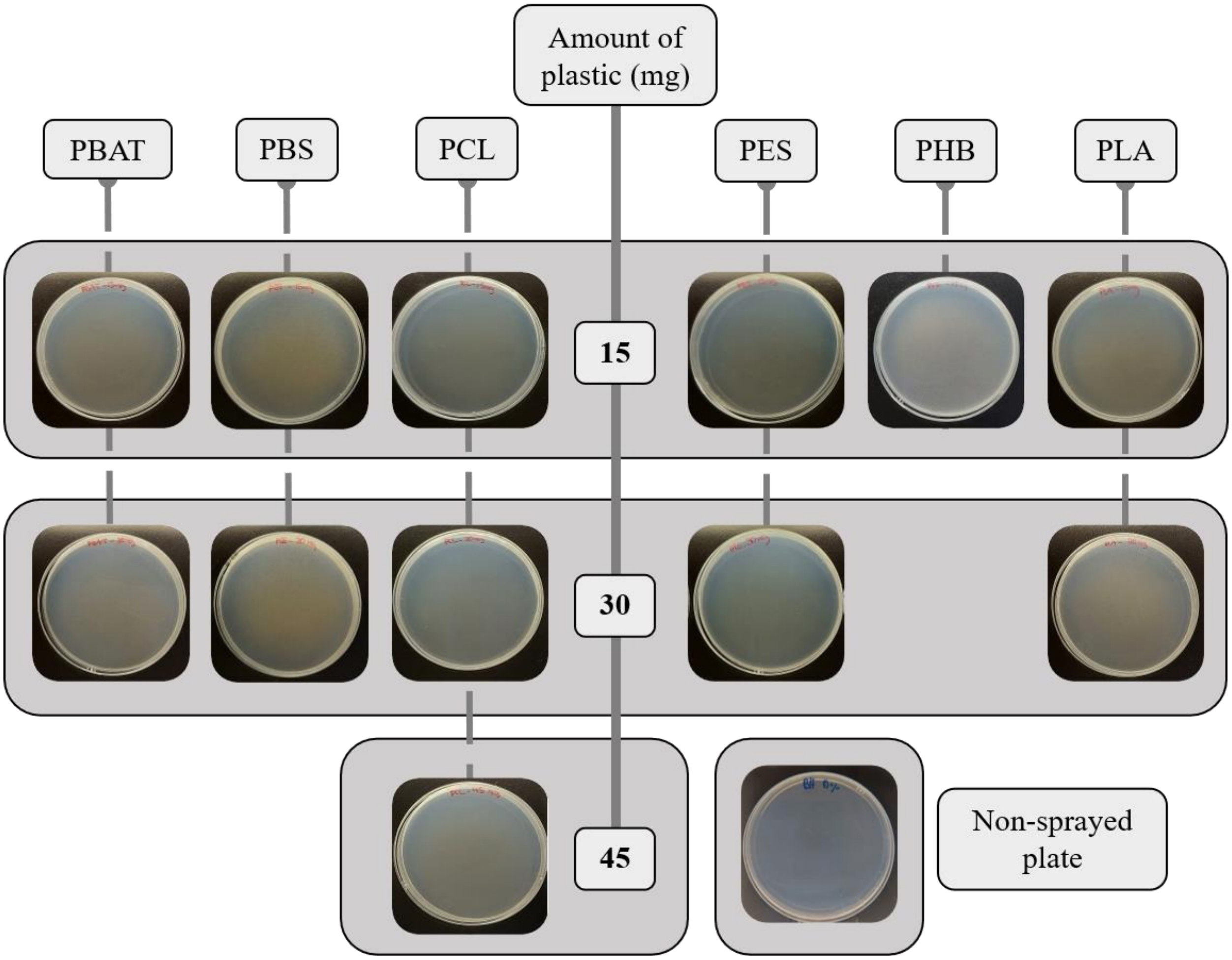


**Figure S2**. Plate opacity of different polyester-sprayed plates depending on the amount of plastic sprayed (15-45 mg). A non-sprayed plate is used as comparison.

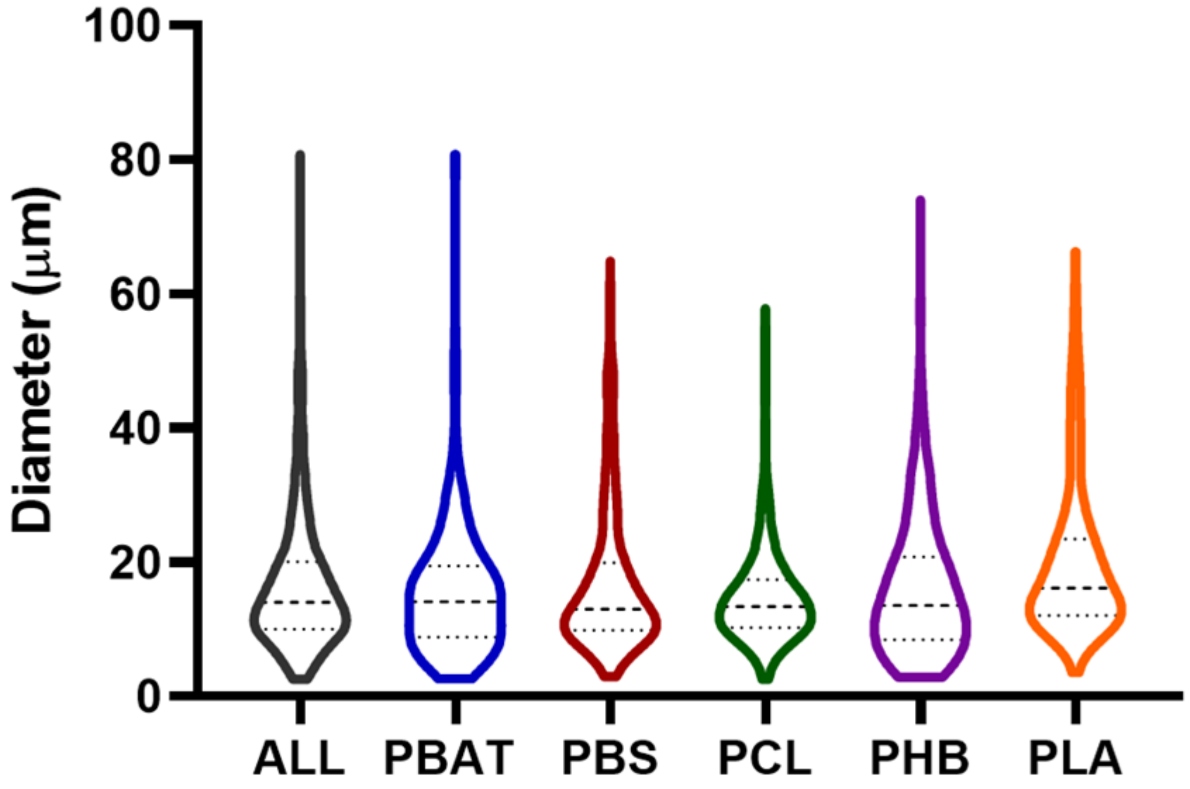


**Figure S3**. Violin plot distribution of polyester microparticles according to their diameter (N = 440 for each individual polyester;). Diameter measurements were made based on optical microscope images.

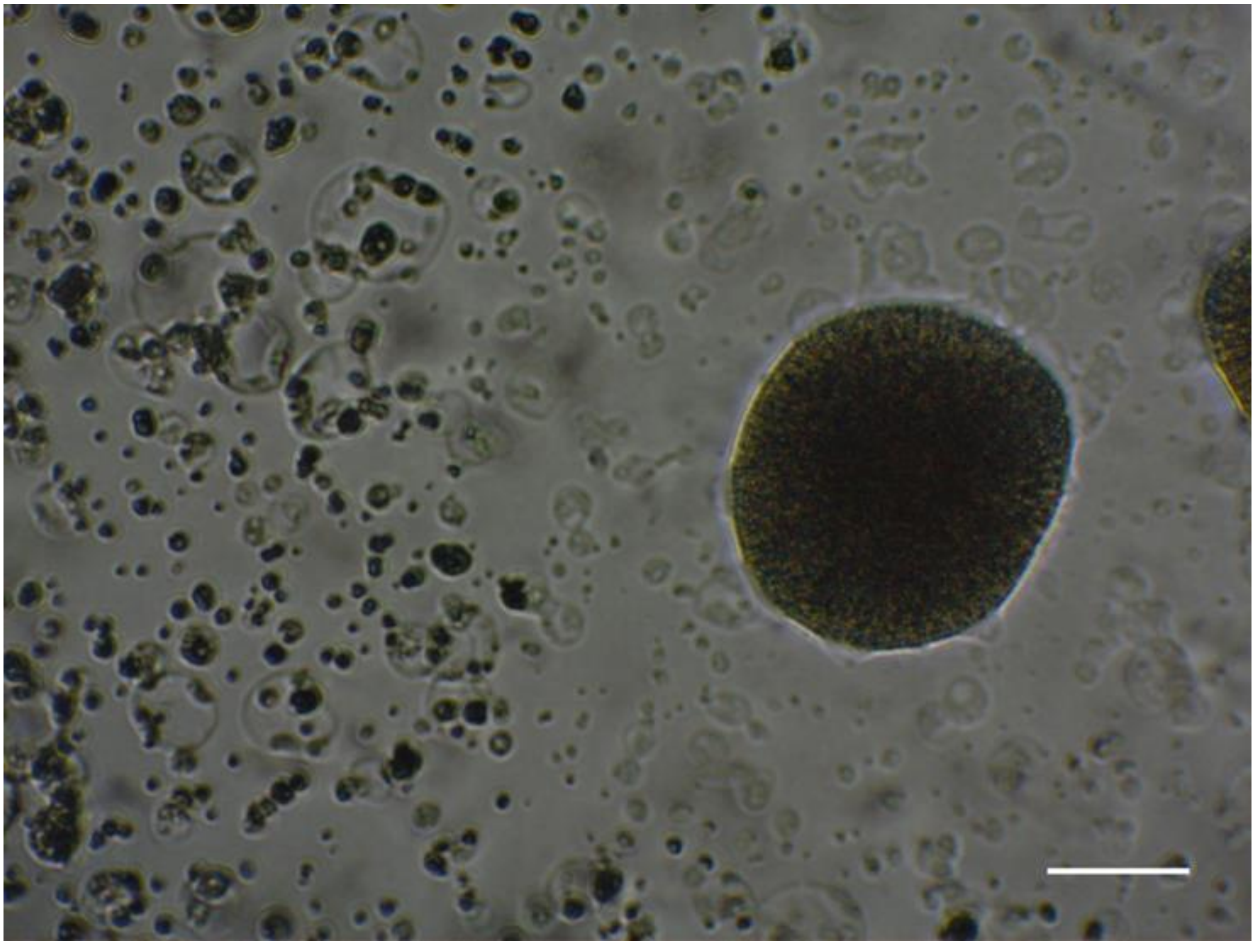

**Figure S4**. PBAT hydrolysis halo around a small colony of *Vishniacozyma* sp. The transparent PBAT microparticles around the colony indicate an early enzymatic depolymerization. Scale bar: 100 µm.

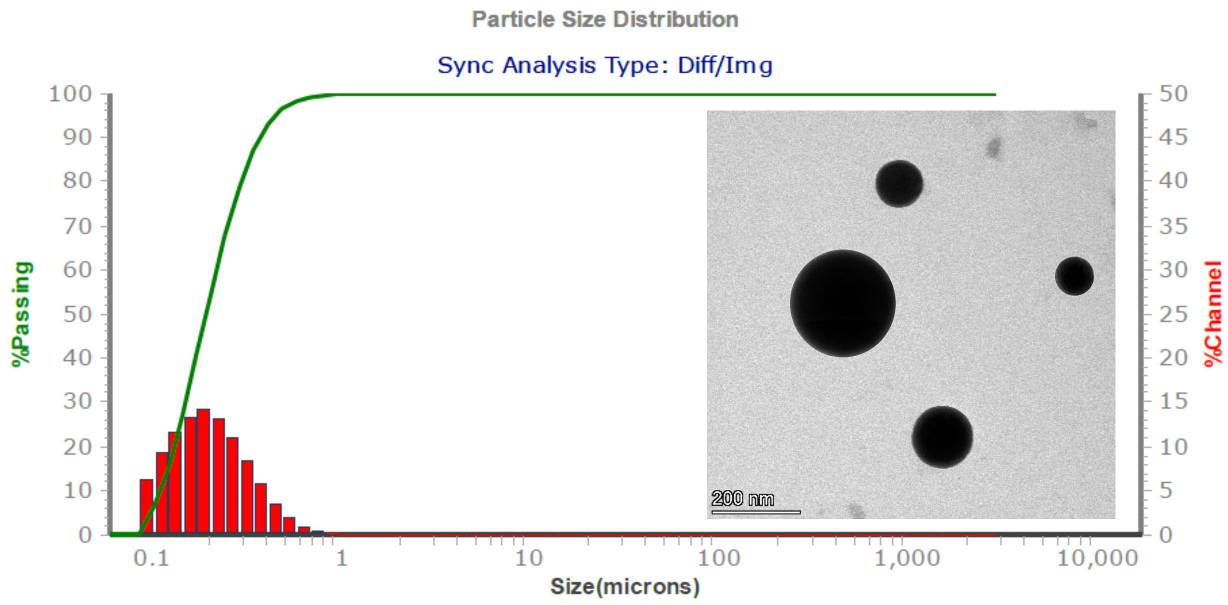


| Median size (µm) | Percentiles (P) (µm) | | Maximum size (µm) | Minimum size (µm) |
|---|---|---|---|---|
| | $P_{20}$ | $P_{80}$ | | |
| 0.193 | 0.131 | 0.295 | 1.635 | 0.102 |

**Figure S5**. Median size of PCL particles obtained from an emulsified solution. Particle size distribution obtained by SLS is shown along with several statistically relevant values. %Passing is the cumulative value of each particle size and %Channel shows the distribution values for each particle size. An HRSTEM image of the same PCL particles is included. Scale bar: 200 µm.

**Table S1**. Descriptive statistical values associated with all polyester microparticle samples.

| | Polyester | | | | | |
|---|---|---|---|---|---|---|
| | All | PBAT | PBS | PCL | PHB | PLA |
| Nº of particles (n) | 2200 | 440 | 440 | 440 | 440 | 440 |
| Median (µm) | 14.04 | 14.10 | 13.02 | 13.40 | 13.55 | 16.16 |
| Q1 (µm) | 10.03 | 8.842 | 9.869 | 10.29 | 8.478 | 12.05 |
| Q3 (µm) | 20.10 | 19.43 | 19.93 | 17.39 | 20.79 | 23.48 |
| Minimum (µm) | 2.590 | 2.645 | 2.958 | 2.590 | 2.857 | 3.541 |
| Maximum (µm) | 80.82 | 80.82 | 64.95 | 57.86 | 74.07 | 66.31 |

**Table S2**. Equations and data used to calculate the estimated SSA values for the air-spraying and emulsification methods.

| | Method | |
|---|---|---|
| | Emulsification | Air-spraying |
| PCL mass/plate ($m_{PCL}$) (g) | 0.025 | 0.015 |
| Particle diameter ($r_P$) (µm) | 0.2 | 14 |
| Particle volume ($V_P$) (µm$^3$) | $\frac{32}{3000}\pi$ | $\frac{1372}{3}\pi$ |
| Particle surface area ($S_P$) (µm$^2$) | $\frac{4}{25}\pi$ | $196\pi$ |
| Total number of particles ($N_P$) | $6.51 \cdot 10^{11}$ | $9.11 \cdot 10^{6}$ |
| Specific surface area (SSA) (µm$^2$/g) | $1.31 \cdot 10^{13}$ | $3.74 \cdot 10^{11}$ |

**<u>List of equations</u>**

$$S_P = 4\pi r_P^{\,2}$$

$$V_P = \frac{4}{3}\pi r_P^{\,3}$$

$$V_{PCL} = \frac{m_{PCL}}{\rho_{PCL}}$$

$$N_P = \frac{V_{PCL}}{V_P}$$

$$SSA = \frac{N_P \cdot S_P}{m_{PCL}}$$

$\rho_{PCL}$ (polycaprolactone density) (25 ºC)= 1.146 g/cm$^3$

PCL volume, $V_{PCL}$
